# Allosteric modulation of dimeric GPR3 by ligands in the dimerization interface

**DOI:** 10.1101/2025.04.19.649678

**Authors:** Zeming Qiu, Wei Wang, Yingying Nie, Junxiang Lin, Beimeng Zhang, Haonan Xing, Sanduo Zheng

## Abstract

Class C G-protein coupled receptors (GPCRs) are widely recognized to function as a dimer, whereas the existence, molecular mechanisms and functional importance of dimerization in class A GPCRs remain poorly explored. GPR3, GPR6 and GPR12 are closely related to cannabinoid receptors, and are associated with neurogenerative diseases such as Parkinson’s and Alzheimer’s diseases. Here, we determined cryo-EM structures of GPR3-G_s_ complex in the active state and GPR3 alone in the inactive state. We observe that GPR3 forms symmetric dimers in both states, with the dimerization interface primarily involving TM5 and TM6. Upon activation, GPR3 undergoes an outward movement of TM6 and an inward movement of TM7, alongside stabilization of helix 8, which is disordered in the inactive state. Functionally, GPR3 dimerization attenuates G_s_ signaling. Furthermore, we identify AF64394 as a negative allosteric modulator that targets the GPR3 dimerization interface and allosterically shifts the α-helical structure of ICL2 into a random coil structure, thereby impairing G_s_ coupling. Our findings provide structural and mechanistic insights into class A GPCR dimerization, offering potential therapeutic strategies targeting the dimerization interface of these receptors.

## Introduction

GPCRs, a largest family of membrane receptors, translate extracellular stimuli into intracellular responses, and play critical roles in various physiological functions and pathological conditions. As a result, GPCRs are the targets of about 36% of all approved drugs (Lorente, Sokolov et al., 2025). Across all species, GPCRs are divided into six main classes (A-F) based on sequence homology (Kolakowski, 1994), and can adopt various oligomerization states (Bouvier, 2001). While Class C and D GPCRs are known to function as dimers both structurally and functionally (Gusach, Garcia-Nafria et al., 2023), the oligomerization of class A GPCRs continues to be a topic of ongoing debate(Chabre & le Maire, 2005, Fotiadis, Jastrzebska et al., 2006). It is now widely accepted that monomeric class A GPCRs is sufficient for engaging and activating G proteins, as evidenced by the fact that cryo-EM structures of the class A GPCR-G protein complex in different intermediate states are determined in a monomeric form (Papasergi-Scott, Perez-Hernandez et al., 2024, Rasmussen, DeVree et al., 2011, Teng, Chen et al., 2022b). However, a growing body of biochemical and biophysical studies suggest that some class A GPCRs can form homodimer or heterodimer (Asher, Geggier et al., 2021, Farran, 2017, Gurevich & Gurevich, 2018, Milligan, Ward et al., 2019). These high-order oligomeric states may play regulatory roles in receptor signaling, trafficking and allosteric modulation. Despite their functional significance, structural evidence supporting the existence of class A GPCR dimerization remains limited.

The transient and weak nature of class A GPCR dimer interactions presents significant challenges for their structural determination using cryo-EM. While X-ray crystal structures of class A GPCRs have identified distinct dimerization interfaces, the physiological relevance of these dimers remains uncertain due to potential crystal packing artifacts. Recently, the cryo-EM structure of the apelin receptor in complex with a G protein was resolved in a dimeric state (Yue, Liu et al., 2022), featuring two receptors coupled to a single G protein. Moreover, the cryo-EM structure of the chemokine receptor CXCR4 alone revealed trimeric and tetrameric states (Saotome, McGoldrick et al., 2025). However, further biochemical and pharmacological studies are needed to confirm the physiological significance of these oligomeric arrangements.

Our previous studies have shown that a subset of class A orphan GPCRs exhibit high constitutive G_s_ signaling (Nie, Qiu et al., 2023). Notably, receptors such as GPR174, GPR3, GPR6, and GPR12 elevate intracellular cAMP levels to an extent comparable to fully ligand-activated receptors, a phenomenon also observed by other research groups (Eggerickx, Denef et al., 1995, Lu, Jang et al., 2021). GPR3/6/12 belong to the same subfamily, and are implicated in various neurological processes including neurite outgrowth (Tanaka, Ishii et al., 2007, Tanaka, Shimada et al., 2022), pain perception (Ruiz-Medina, Ledent et al., 2011), mood addition (Valverde, Celerier et al., 2009) and motor control (Oeckl, Hengerer et al., 2014). In the nervous system, GPR3 promotes amyloid-beta production through β-arrestin recruitment by modulating γ-secretase activity (Huang, Rafael Guimaraes et al., 2022, Huang, Skwarek-Maruszewska et al., 2015, Thathiah, Spittaels et al., 2009), making it a potential drug target for Alzheimer’s disease. GPR6 is restrictively expressed in the indirect pathway D2-type medium spiny neurons (MSN), counteracting D2 dopamine receptor signaling with respect to cAMP production. It could affect the output of the indirect pathway, preventing involuntary movements (Oeckl et al., 2014). A selective inverse agonist of GPR6, CVN424 has entered Phase 3 clinical trial for treatment of Parkison’s disease, offering a non-dopaminergic approach to avoid dyskinesia. In the reproductive system, GPR3 and GPR12 maintain meiotic arrest in oocytes before ovulation by sustaining cAMP levels (Freudzon, Norris et al., 2005, Hinckley, Vaccari et al., 2005, Ledent, Demeestere et al., 2005, Mehlmann, Saeki et al., 2004). Moreover, lipolysis-driven expression of GPR3 in adipose tissue promotes heat production and energy expenditure, preventing mice from obesity (Dong, Hu et al., 2024, Godlewski, Jourdan et al., 2015, Sveidahl Johansen, Ma et al., 2021).

During our deorphanization efforts, we unexpectedly discovered that GPR3, unlike GPR6 and GPR12, functions as a homodimer, as evidenced by cryo-EM structures of both the GPR3-G_s_ complex and GPR3 alone. We further validate the physiological relevance of GPR3 dimer by identifying a negative allosteric modulator that specifically targets this dimerization interface.

## Results

### Dimeric assembly of GPR3 in the active state

Building on our previous discovery that the endogenous ligand lysoPS binds to GPR174, resulting in its high constitutive activity (Nie et al., 2023), we hypothesized that the high constitutive activities of GPR3, GPR6, and GPR12 are likely due to the binding of endogenous ligands. To investigate the identity of these ligands, we aimed to determine the cryo-EM structure of the GPR3-G_s_ complex. To improve the stability of the GPR3-G_s_ complex, we introduced mutations at three residues (A^34.50^L^34.51^T^34.52^) in the ICL2 of GPR3, replacing them with the corresponding residues (P^34.50^F^34.51^K^34.52^) from the β2 adrenergic receptor (β2AR) (Rasmussen et al., 2011) (referred to as GPR3PFK), which contains bulky hydrophobic side chains known to play a crucial role in G_s_ coupling (**Figure 1-figure supplement 1A**).

**Figure 1.**
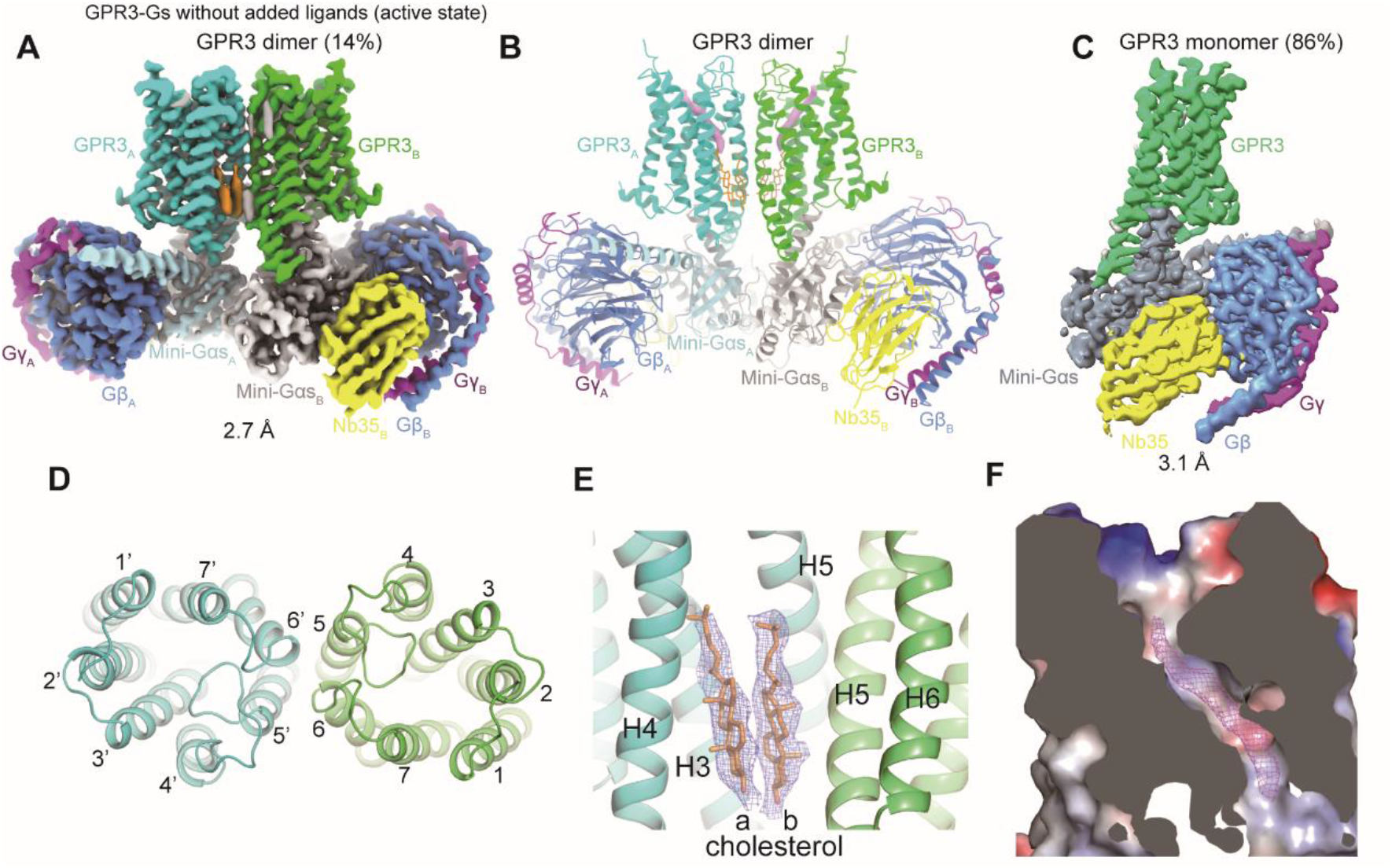
GPR3 forms a dimer in its active state. (**A**) Cryo-EM map of the dimeric GPR3-Gs complex, showing the GPR3 dimer (cyan, green), two copies of mini-Gαs (light blue, grey), Gβ (blue), Gγ (magenta), and Nb35 (yellow). (**B**) Overall structure of the dimeric GPR3-Gs complex, colored according to the scheme in (**A**). (**C**) Cryo-EM map of the monomeric GPR3-Gs complex. (**D**) Extracellular view of the GPR3 dimer structure. (**E**) EM density showing the two parallel cholesterol molecules at the dimerization interface of GPR3.

(**F**) Electrostatic potential map of the potential endogenous lipid-binding surface, with the corresponding EM density for the lipid shown.

Indeed, GPR3PFK shows higher basal activity than the wild-type (WT), suggesting improved G_s_ coupling (**Figure 1-figure supplement 1B, C**). Following our established protocols (Nie et al., 2023), we purified GPR3PFK and mini-Gαs fusion protein, assembling it with purified Gβγ and Nb35 (**Figure 1-figure supplement 1D**). Surprisingly, cryo-EM analysis of GPR3PFK-G_s_ complex revealed the presence of both dimeric (14%) and monomeric (86%) assemblies, with respective overall resolution of 2.7 and 3.1 Å (**Figure 1A-C, and Figure 1-figure supplement 1F-I and Table 1**). Monomeric GPR3 exhibits a characteristic arrangement of active GPCRs coupled to G_s_ (**Figure 1C**). While dimeric GPR3 engages two copies of mini-Gαs lacking the AHD domain, the presence of two full-length Gαs molecules would result in a structural clash between their AHD domains, making such an arrangement incompatible (**Figure 1-figure supplement 1J**). This also explains why the dimeric GPR3 is not observed in recent published structures of the GPR3-G_s_ complex that use full-length Gαs (Chen, Staffen et al., 2024, Russell, Zhang et al., 2024, Xiong, Xu et al., 2024). Therefore, dimeric GPR3 likely recruit only one G_s_ heterotrimer, akin to glutamate receptors that can activate a single G protein despite their dimeric assembly. Notably, compared to the apelin receptor (207 Å^2^) (Yue et al., 2022), the dimerization interface of GPR3 buries a larger surface area (839 Å^2^), primarily involving TM5 and TM6 (**Figure 1D**), suggesting the physiological relevance of the dimer. Moreover, two parallel cholesterol molecules with well-defined EM density are localized at the dimerization interface, possibly stabilizing the dimer (**Figure 1E**). Reminiscence of GPR174, the structure reveals a well-defined density for the putative endogenous ligands bound in the extended hydrophobic groove (**Figure 1F**), a feature also observed in recently published structures of GPR3-G_s_ (Chen et al., 2024, Russell et al., 2024, Xiong et al., 2024) and GPR6-G_s_ (Barekatain, Johansson et al., 2024) complexes. The ligand is likely derived from cells or culture medium, and copurified with the receptors due to their high affinity binding.

### Key polar residues of GPR3/6/12 involved in endogenous ligand binding

The copurified ligand is encircled by a cluster of hydrophobic residues, suggesting that it likely contains an extended, unbranched alkyl group originating from a lipid (**Figure 2A**). Two polar residues H96^2.60^ and Y280^7.36^ are positioned near the lipid, and may form hydrogen bonds with its polar moiety. All seven TMs from the extracellular portion are involved in binding the lipid, accounting for the high affinity binding. To determine if the bound lipid contributes to the high constitutive activity, we conducted Glosensor cAMP accumulation assay to evaluate the effect of mutations in these residues on basal activity. Interestingly, when GPR3 was expressed at high levels using the human crytomegalovirus (CMV) promoter, none of the mutations affected the basal activity (**Figure 2B**). However, expressed at low levels using the spleen focus-forming virus (SFFV) promoter, mutations of H96^2.60^ or Y280^7.36^ of GPR3 dramatically impaired basal activity, compared to mutations in other hydrophobic residues such as L116^6.32^, F120^3.36^, I124^3.40^, and F202^5.47^, which are involved in binding the lipid’s alkyl moiety. Similarly, mutations of H128^2.60^ or Y312^7.36^ as well as other hydrophobic residues significantly impaired the basal activity of GPR6 at low levels, with double mutations of the two polar residues almost abolishing the basal activity of GPR6 even at high expression levels (**Figure 2C**). The discrepant results between high and low expression levels are expected, as high receptor expression often increases apparent potency of ligands. As a result, the concentration of the endogenous lipid produced by cells is above its concentration that is required to achieve maximal response in GPR3 or GPR6 mutants at high expression levels due to the lipid’s higher potency, whereas at low expression levels, its lower potency prevents the mutants from reaching a maximal response, leading to reduced basal activity. In contrast to GPR3 and GPR6, mutation of the equivalent residues N100^2.60^ or Y284^7.36^ in GPR12 abolished its basal activity at high expression level, likely due to the lower potency of endogenous lipid for GPR12 compared to GPR3 or GPR6 (**Figure 2D**). The expression levels of all mutants were adjusted to be comparable to WT, ensuring that differences in basal activity were not influenced by expression levels (**Figure 2-figure supplement 1A-E**). Although previous studies proposed oleic acid (Xiong et al., 2024), oleoylethanolamide (Chen et al., 2024), sphingosine 1-phosphate (Uhlenbrock, Gassenhuber et al., 2002) and sphingosylphosphorylcholine (Ignatov, Lintzel et al., 2003) as putative ligands of GPR3, none of these lipids can activate GPR3 in our cAMP accumulation assay (**Figure 2-figure supplement 1F-H**). In addition, we tested other lipids including cannabinoids (**Figure 2-figure supplement 1F**), fatty acids with varying length of acyl chain (**Figure 2-figure supplement 1G**), and lysolipids/sphingolipids (**Figure 2-figure supplement 1H**), none of which show any activity. Our previous studies have shown that GPR174 cannot be activated by exogenous lysophosphatidylserine (lysoPS) because of its maximal activation by endogenous lysoPS produced by cells (Nie et al., 2023). Similarly, endogenous lipids may maximally activate GPR3/6/12, preventing them from responding to exogenous lipids. Therefore, we cannot rule out the possibility that one of these test lipids is the endogenous ligand of GPR3/6/12. The search for their endogenous ligands is ongoing in our laboratory. In summary, these findings suggest that endogenous lipids induce a maximal response in GPR3/6/12, leading to high basal activity, and that two polar residues are critical for binding the polar moiety of the endogenous lipid.

**Figure 2.**
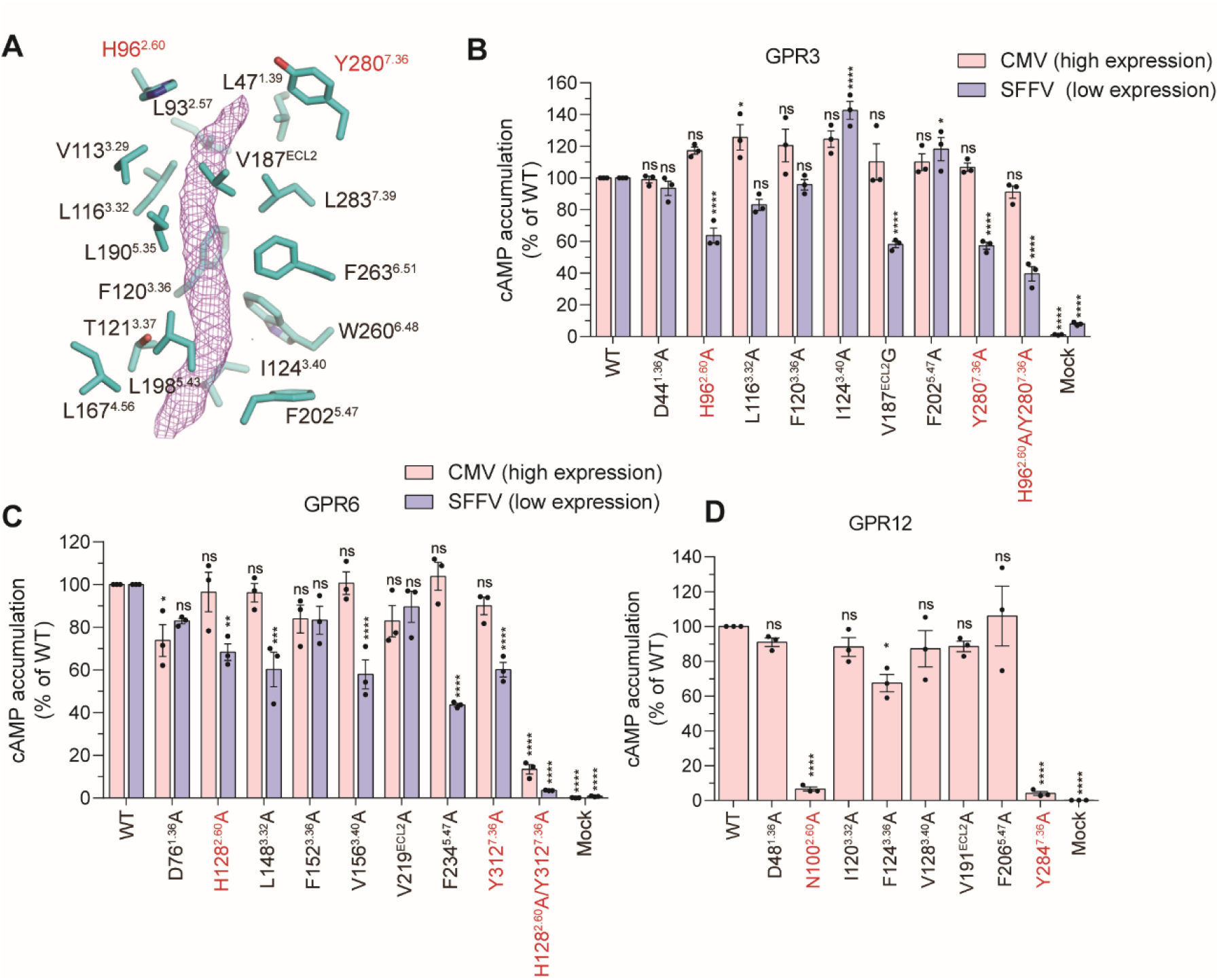
Putative endogenous lipid-binding site. (**A**) GPR3 residues in close proximity to the EM density corresponding to endogenous lipids are shown as stick representations. Two polar residues, H96^2.60^ and Y280^7.36^ are labelled in red. (**B-C**) Effect of mutations in the putative lipid-binding site on cAMP levels in cells expressing GPR3 (**B**) or GPR6 (**C**) mutants, driven by the CMV and SFFV promoters. (**D**) Effect of mutations in the putative lipid-binding site of GPR12 on cAMP levels in cells expressing GPR12 mutants under the CMV promoter. All cAMP levels were measured using the cAMP accumulation assay and normalized as a percentage of the wild-type (WT) levels.

### Dimeric GPR3 in the inactive state and mechanism of GPR3 activation

To exclude the possibility that dimeric GPR3 in the active state results from the use of mini-G_s_ or mutations in GPR3, we aimed to determine the cryo-EM structure of GPR3 alone in the inactive state. To facilitate particle alignment of GPR3, we employed a GPCR fusion strategy where the intracellular loop 3 (ICL3) is replaced by a thermostabilized apocytochrome b562 (BRIL) that is combined with an anti-BRIL Fab fragment and an anti-Fab nanobody (**Figure figure supplement 1A**)(Tsutsumi, Mukherjee et al., 2020, Zhang, Wu et al., 2022). Cryo-EM analysis of the GPR3-BRIL fusion protein also revealed both monomeric and dimeric GPR3, with the dimeric form being more predominant (**Figure 3-figure supplement 1B, C**). The structure of dimeric GPR3 was determined at a global resolution of 3.32 Å (**Figure 3-figure supplement 1D-F and Table 1**). The dimer is not symmetrical since one copy of the BRIL-Fab-Nb module is more flexible than the other one (**Figure 3A, B**). Interestingly, a lipid-like EM density is also observed in the inactive GPR3 structure, but it appeared less defined compared to the active GPR3 structure (**Figure 3C**), suggesting that G protein binding further stabilize the endogenous lipid binding to GPR3. Moreover, one cholesterol located at the dimerization interface near the endogenous lipid exhibits poorly-defined EM density, indicating a potential cooperative binding between this cholesterol and the lipid (**Figure 3A**). While activation of dimeric class C GPCRs involves structure rearrangement of the dimerization interface in the transmembrane (TM) helices, the active GPR3 share same dimerization interface with the inactive structure. The structure of the GPR3 dimer in its active state closely resembles that of the GPR3 monomer in the active state (**Figure 3D**). Comparing the active and inactive states of the dimeric GPR3 reveals conformational changes linked to receptor activation, which are consistent with those observed in most class A GPCR activation. Lipid binding causes a slight compaction of the orthosteric binding pocket, driven by the inward movement of TM1, TM2, and TM7 (**Figure 3E**). Notably, H96^2.60^ and Y280^7.36^, which are crucial for receptor activation, shift toward the lipid due to their interaction. Furthermore, lipid binding triggers a downward shift of the toggle switch W260^6.48^, leading to the outward movement TM6 (**Figure 3F**). Y297^7.53^, part of the conserved motif NP^7.50^xxY in TM7 moves toward the position previously occupied by TM6 in the inactive state, potentially forming a water-mediated hydrogen bond with Y213^5.58^. These movements result in the inward shift of TM7 and a slight movement of TM5 toward TM6 (**Figure 3G**). The C-terminal helix 8 becomes more rigid as a result of the inward movement of TM7. In most class A GPCRs such as β2AR, R^3.50^ of the conserved D(E)^3.49^R^3.50^Y^3.51^ motif forms an ionic lock with E^6.30^, stabilizing the inactive state. However, in GPR3, E^6.30^ is replaced by T248^6.30^, which may contribute to its high basal activity (**Figure 3G**).

**Figure 3.**
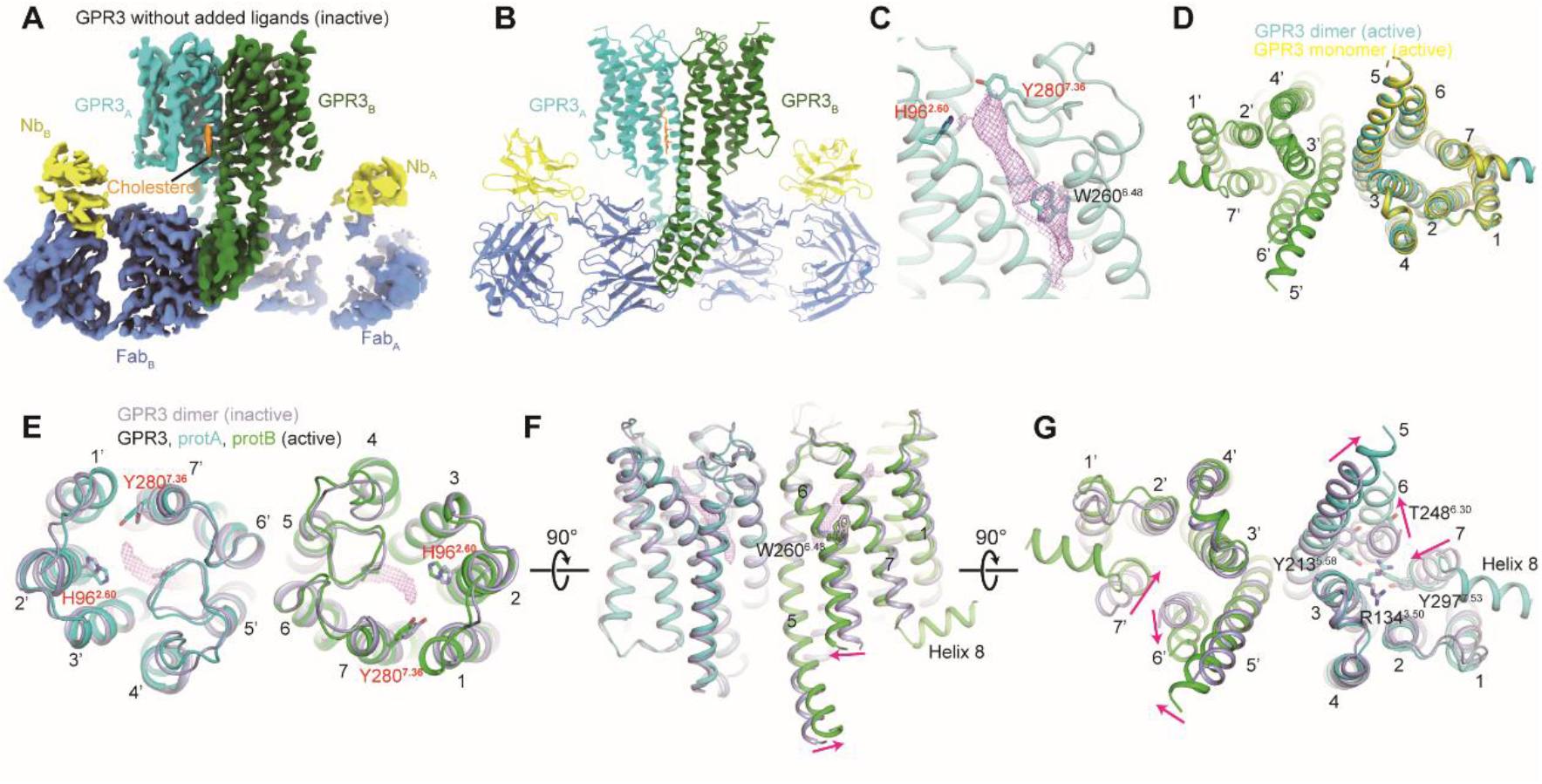
GPR3 forms a dimer in its inactive state. (**A**) Cryo-EM map of the GPR3-BRIL fusion protein (cyan, green) in complex with anti-BRIL Fab (blue) and anti-Fab nanobody (yellow). (**B**) Overall structure of the GPR3-BRIL fusion complex, using the same color scheme as in (A). (**C**) EM density map of the endogenous lipid (shown in magenta mesh), with key polar residues H96^2.60^, Y280^7.36^, and the toggle switch W260^6.48^ represented as sticks. (**D**) Structural overlay of the active GPR3 monomer (yellow) with a protomer (cyan) of the GPR3 dimer. (**E-G**) Superposition of dimeric GPR3 structures in the inactive and active states, shown from the extracellular view (**E**), side view (**F**), and cytoplasmic view (**G**).

### Functional significance of the dimerization

Like class C GPCRs, the dimerization interface is compact and extensive, primarily formed by TM5 and TM6 from both protomers. On the extracellular side of the TMs, hydrophobic interactions play a crucial role in stabilizing the interface. Key residues contributing to these interactions include V196^5.41^, I200^5.45^, and M204^5.49^ on TM5 of one protomer, and F203^5.48^, F206^5.51^, V265^6.53^, L268^6.56^, and L269^6.57^ on TM5 and TM6 of the other protomer (**Figure 4A**). This interface is further stabilized by a salt bridge interaction between K192^5.37^ and D271^ECL3^. On the intracellular side, a network of hydrogen bonds reinforces the interface. Specifically, Q211^5.56^ on one protomer forms a hydrogen bond with its counterpart in the other protomer. Furthermore, Y135^3.51^ of the DRY motif in one protomer is stabilized by hydrogen bonds with Q215^5.60^ from both protomers. In contrast, Q215^5.60^ is substituted by an R in the equivalent position of many class A GPCRs including GPR6 (**Figure 4A and Figure 4-figure supplement 1**), which likely destabilizes this interface. Moreover, several hydrophobic residues of GPR3 involved in stabilizing the interface is replaced by residues with shorter hydrophobic side chains such as A or V in GPR6. These substitutions prevent GPR6 from forming a stable dimerization interface. Indeed, cryo-EM analysis of the GPR6-BRIL fusion protein reveals a monomeric form as well as an antiparallel GPR6 dimer that is not compatible with the membrane, indicating that GPR6 can only form a monomer in solution (**Figure 4B**). To investigate the functional significance of GPR3 dimerization, we created a monomeric GPR3 by replacing the residues in the dimerization interface with their counterparts in GPR6 (referred to as the GPR3_6 chimera). As expected, fluorescence size-exclusion chromatography (FSEC) revealed that the GPR3_6 chimera exists in a monomeric form, whereas both monomeric and dimeric forms are present in GPR3WT (**Figure 4C**). We next assessed the basal activity of the GPR3_6 chimera toward G_s_ signaling using the cAMP accumulation assay. The chimera exhibited enhanced G_s_ coupling at both high and low expression levels (**Figure 4D**). This observation can be attributed to the structural constraint of the dimeric GPR3, in which only one protomer engages with the G protein while the other remains unbound, thereby limiting the overall coupling efficiency compared to the monomeric form.

**Figure 4.**
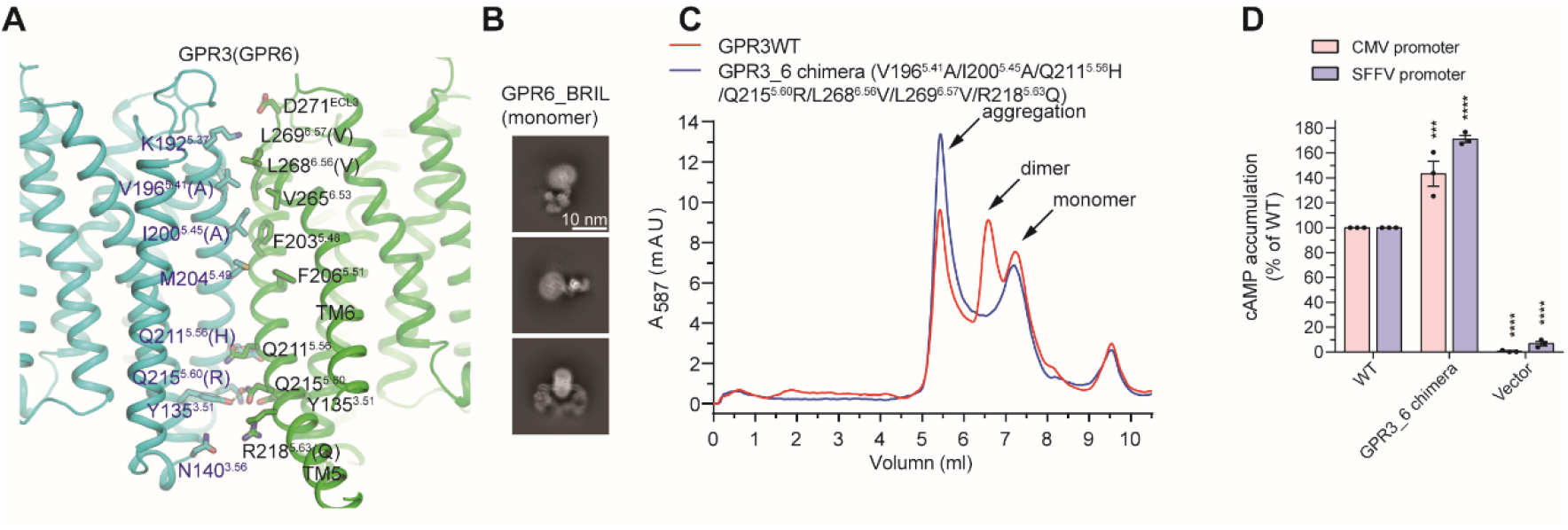
Dimerization interface of GPR3. (**A**) Residues involved in GPR3 dimer formation are shown as sticks. Non-conserved residues at the dimer interface in GPR6 are indicated in parentheses. (**B**) 2D class averages of the GPR6-BRIL fusion protein in complex with anti-BRIL Fab and an anti-Fab nanobody. (**C**) Fluorescence size-exclusion chromatography profiles of GPR3WT and the GPR3_6 chimera, in which dimer interface residues in GPR3 are replaced with the corresponding residues from GPR6. (**D**) cAMP levels in Expi293F cells expressing GPR3WT or the GPR3_6 chimera, driven by CMV and SFFV promoters, measured using the GloSensor cAMP accumulation assay.

### Allosteric modulation of GPR3 by AF64394 in the dimerization interface

Previous studies identified a selective GPR3 inverse agonist AF64394. Indeed, AF64394 selectively reduced the basal activity of GPR3 in a dose-dependent manner, with an IC50 of 42.9 nM (**Table 2**), while showing no activity on GPR6, GPR12 or fully activated D1 dopamine receptor (D1R) (**Figure 5A**). To investigate molecular mechanisms underlying inverse agonism of AF64394, we determined the cryo-EM structure of GPR3-BRIL bound to AF64394 (**Figure 5B and Figure 5-figure supplement 1 and Table 1**). Unlike the structure of GPR3-BRIL alone, the presence of AF64394 causes GPR3 to form only a dimer, with no monomer formation detected (**Figure 5-figure supplement 1C**), suggesting that it stabilizes the GPR3 dimer. A well-defined density corresponding to AF64394 was observed at the dimerization interface of GPR3, while a separate density for endogenous lipids was detected in the orthosteric binding pocket (**Figure 5C, D**), indicating that AF64394 does not interfere with agonist binding. Notably, AF64394 is situated in a hydrophobic groove formed by TM3, TM4 and TM5 of one protomer in the cytoplasmic side. Residues mediating hydrophobic interactions with AF64394 include L128^3.44^, V132^3.48^, R152^4.41^, V155^4.44^, M156^4.45^, L159^4.48^, I208^5.53^ and L212^5.57^. In contrast, only two residues, R218^5.63^ and Q211^5.66^ on TM5 of the other protomer contribute to hydrophilic interactions with the pyrimidine ring and its attached nitrogen group, respectively (**Figure 5D**). Notably, R218^5.63^ in GPR3 is substituted by Q in GPR6 and K in GPR12, explaining AF64394’s selective binding with GPR3 (**Figure 4-figure supplement 1**). Consistent with our structural findings, mutations of H96 or Y280 in the orthosteric binding pocket had minimal effect on the activity of AF64394, whereas mutations of R218^5.63^ and Q211^5.66^ in one protomer almost completely abolished its activity (**Figure 5E and Table 2**). Mutations of hydrophobic residues in the other protomer reduced the potency of AF64394 to varying degree, with R152^4.41^ and V132^3.48^ showing the most pronounced effect.

**Figure 5.**
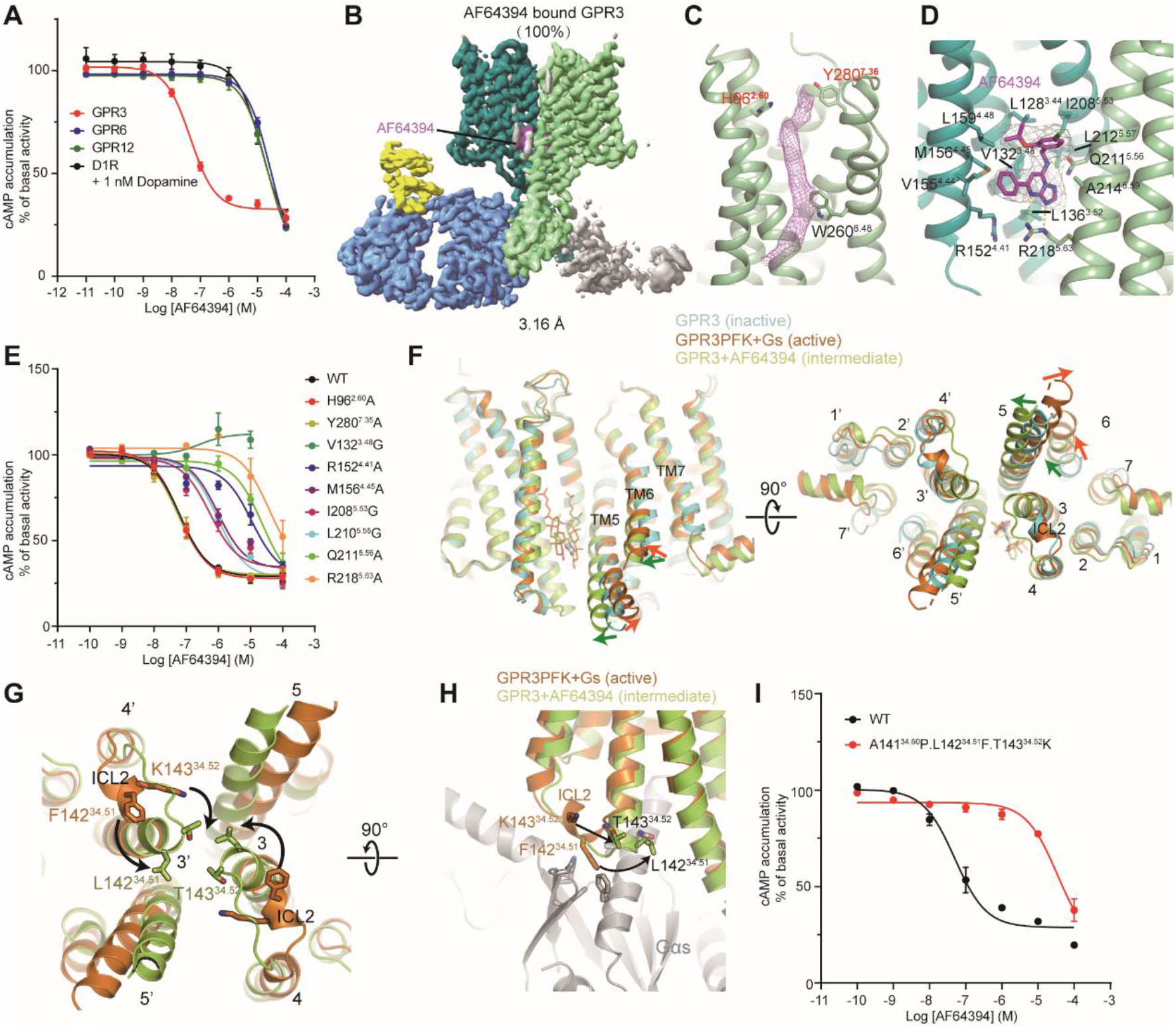
Structural and functional characterization of AF64394 binding to GPR3. (**A**) cAMP accumulation assay showing the effect of AF64394 on GPR3, GPR6, GPR12, and D1R in the presence of 1 nM dopamine. (**B**) Cryo-EM map of AF64394-bound GPR3, highlighting the ligand (magenta), GPR3-BRIL (green and cyan), anti-BRIL Fab (blue) and an anti-Fab nanobody (yellow). The Fab and nanobody are poorly defined in one protomer, with the corresponding map colored in grey. (**C**) EM density of the endogenous lipid shown in magenta mesh, with key interacting residues H96^2.60^, Y280^7.36^, and W260^6.48^ represented as sticks. (**D**) Close-up view of the AF64394 binding pocket. The EM density map for AF64394 is shown in grey mesh. Yellow dash line indicates hydrogen bonds. (**E**) cAMP accumulation assay in Expi293F cells expressing WT or mutant GPR3, showing the effect of site-specific mutations on the AF64394 response. (**F**) Superposition of inactive GPR3, active GPR3PFK, and AF64394-bound GPR3 intermediate structures, revealing conformational changes upon AF64394 binding. (**G**) Comparison of the active state and the AF64394-bound state of GPR3, showing a shift in ICL2 from an α-helical to a random coil structure. (**H**) Close-up view of interactions between the α-helical structure of ICL2 with Gαs in the active state. The transition of ICL2 to a random coil structure upon AF64394 binding disrupts G_s_ coupling. (**I**) cAMP accumulation assay in Expi293F cells expressing WT or triple-mutant GPR3PFK, demonstrating altered AF64394 response.

AF64394 occupies the same position as the two parallel cholesterol molecules observed in the active structure (**Figure 5F**). Upon ligand binding, the extracellular portions of the TMs in GPR3 remain largely unchanged, whereas the intracellular portions of TMs, particularly TM5, TM6 and TM7 from both protomers, shift toward AF64394. This movement results in a narrower pocket at the dimerization interface. Despite the outward movement of TM6 and inward shift of TM7, the conformation of GPR3 bound to AF64394, referred to as an intermediate state, differs from the active state due to the opposing shift of TM5 (**Figure 5F**). In the active state, ICL2 adopts an α-helical conformation, allowing L142^34.51^ (F142^34.51^ in GPR3PFK) to insert into a hydrophobic pocket on Gαs (**Figure 5G, H**). A mutation of L142^34.51^ to alanine completely abolishes the basal activity of GPR3, highlighting its critical role in G_s_ coupling (**Figure 1-figure supplement 1C**). Upon AF64394 binding, however, the α-helical structure of ICL2 transitions into a random coiled structure, driven by hydrophobic interactions between L142^34.51^ and T143^34.52^ from both protomers (**Figure 5G, H**). In the GPR3PFK mutant, a A141^34.50^L142^34.51^T143^34.52^ to PFK mutation likely prevents the conformational transitions of ICL2 from an α-helical to a random coiled structure. This is due to steric clashes between F142^34.51^ and K143^34.52^, which explains the inability of to suppress the G_s_ coupling of the GPR3PFK mutant (**Figure 5I**). Collectively, these results indicate that AF64394 allosterically stabilizes the ICL2 of GPR3 in a random coil structure by targeting the dimerization interface, thereby preventing G_s_ coupling.

## Discussion

There is ongoing debate about whether class A GPCRs can function as an oligomer. Here, we show that GPR3 can form a homodimer in both active and inactive states. In contrast to the class D Ste2 dimer that engages two G proteins (Velazhahan, Ma et al., 2021), dimeric GPR3 can recruit only one G_s_ because of potential steric clashes between two G_s_ when both are simultaneously coupled by GPR3. Therefore, disrupting the dimerization interface enhances G_s_ coupling efficiency, consistent with recent structural studies on the dimeric apelin receptor (Yue et al., 2022). Interestingly, GPR3 features a distinct and more extensive dimerization interface compared to the apelin receptor, suggesting that dimerization interface in class A GPCRs is not conserved. During receptor activation process, the dimerization interface of GPR3 remains unchanged, whereas activation of class C GPCRs including GABA_B_ (Shen, Mao et al., 2021) and glutamate receptors (Lin, Han et al., 2021) involves structural rearrangement at the dimerization interface. In addition, unlike class C GPCRs, which are obligate dimers, GPR3 and the apelin receptor exist in a dynamic equilibrium between monomeric and dimeric states. This equilibrium suggests that dimerization in class A GPCRs may act as a desensitization mechanism, reducing G protein signaling when receptor expression levels are high and dimer formation is favored. Moreover, further studies are required to determine whether the dimerization in GPR3 or the apelin receptor affects ligand pharmacology and the recruitment of other downstream effectors.

Several lipids have been proposed as endogenous ligands of GPR3. Our studies showed that none of lipids activate GPR3 using the GloSensor cAMP accumulation assay. Maximal activation of GPR3 by endogenous lipids produced by cells prevents its responsive to exogenous ligands, making identification of endogenous ligands challenging. Noteworthily, variations in lipid concentrations across different cell lines, along with differences in ligand potency across various signaling assays, may lead to inconsistent results for these receptors, when tested using different assays or different cell lines. For example, GPR174 cannot be activated by exogenous lysoPS using the cAMP accumulation assay, because the cellular lysoPS concentration exceeds the level required for the maximal response (Nie et al., 2023). In contrast, when using the NanoBiT G_s_ recruitment assay (Nie et al., 2023) or the TGFα shedding assay (Inoue, Ishiguro et al., 2012), GPR174 exhibits a dose-dependent manner by lysoPS. This activation occurs because the cellular lysoPS concentration is lower than the potencies using these assays. Based on these findings, we propose three criteria for the endogenous lipid of GPR3/6/12: (I) The lipid must be abundantly present in various cell types, as these receptors exhibit high basal activity across different cell lines; (II) The lipid should have nanomolar potency, ensuring that endogenous production is sufficient for maximal receptor activation; (III)Two key polar residues, H/N^2.60^ or Y^7.36^, must play a crucial role in lipid binding. More rigorous studies are needed to identify the endogenous lipid by depleting it from cells or removing it from the receptors.

Some allosteric modulators act through the dimerization interface of the class C GPCRs in the TMs, suggesting that the dimerization interface is druggable (Shen et al., 2021). Ligands targeting the dimerization interface may exhibit unique pharmacological properties. For GPR3, the endogenous lipid is likely abundant in vivo, and binds with GPR3 with high affinity, making it difficult to develop a potent orthosteric antagonist that can effectively compete with the endogenous lipid. The negative allosteric modulator, AF64394, located at the dimerization interface reduces the basal activity of GPR3 by inducing its conformational changes without disrupting lipid binding. Interestingly, AF64394 promotes the outward movement of TM5 and TM6 while shifting TM7 inward. This is distinct from the active state where the TM5 moves in the opposite direction. These conformation rearrangements cause the α-helical structure of ICL2 to shift to a random coil structure, thereby impairing G_s_ coupling. Conversely, the previously reported positive allosteric modulators, Cmpd-6 and LY315402 stabilize the α-helical structure of ICL2 in β2AR (Liu, Masoudi et al., 2019) and D1R (Teng, Chen et al., 2022a, Zhuang, Krumm et al., 2021), respectively. Furthermore, AF64394 can stabilize the GPR3 dimer. This suggests that, even for GPCRs that that do not typically form dimers or form only transient dimers, it is possible to develop molecular glues that promote dimer formation and allosterically modulate receptor pharmacology. This study provides a framework for developing allosteric modulators that target the dimerization interface of class A GPCRs to achieve more diverse pharmacological properties.

## Methods

### Cloning

The coding sequences of GPR3, GPR6, and GPR12 were cloned from the PRESTO-Tango GPCR Kit (Kroeze, Sassano et al., 2015). Each construct included an N-terminal HA signal peptide (MKTIIALSYIFCLVFA) followed by a Flag tag (DYKDDDDK). For expression under a strong promoter, the constructs were cloned into the pcDNA3.1 vector with the cytomegalovirus (CMV) promoter. For lower expression, the SFFV promoter replaced the CMV enhancer and promoter in the pcDNA3.1 backbone.

For the GPR3–miniGα_s_ fusion construct, GPR3 was fused to miniGα_s_399 (Carpenter & Tate, 2016) via a flexible 34-amino-acid glycine-serine (GS) linker. Three point mutations—A141P, L142F, and T143K—were introduced into the GPR3 coding region using the QuikChange site-directed mutagenesis method following standard protocols. For the GPR3-BRIL and GPR6-BRIL fusion constructs, the ICL3 of GPR3 (residues 223-239) and GPR6 (residues 255-271) was replaced by the BRIL using the linker sequences derived from the A2A-BRIL fusion protein structure (PDB: 4EIY). To potentially stabilize the receptors in the inactive state by restricting the conformational flexibility of TM6, the mutations T248C/R300C (GPR3) and T280C/R332C (GPR6) were introduced.

### Expression and purification of Gβ_1_γ_2_-C68S, Nb35, BAG2 (anti-BRIL-Fab) and anti-Fab-Nb

The coding sequences for the 8×His-tagged heavy and light chains of BAG2, an anti-BRIL Fab, were cloned into the pETDuet-1 vector and transformed into *Escherichia coli* (*E. coli*) BL21 (DE3) cells. A total of 4 L of bacterial culture was grown in LB medium at 37 °C until the OD600 reached 0.6, followed by induction with 200 μM isopropyl β-D-1-thiogalactopyranoside (IPTG). The culture was then shaken at 16 °C for 16 hours. Cells were harvested by centrifugation, resuspended in NTA Buffer (20 mM HEPES, pH 7.4; 150 mM NaCl; 20 mM imidazole), and lysed using a French press high-pressure cell disruptor. The lysate was clarified by centrifugation at 35,000 × g for 1 hour at 4 °C, and the supernatant was filtered through a 0.45 μm membrane filter before being loaded onto Ni-NTA resin via gravity flow. The resin was washed with the NTA buffer, and BAG2 Fab was eluted using NTA Elution Buffer (20 mM HEPES, pH 7.4; 150 mM NaCl; 250 mM imidazole). The protein was further purified using a Superdex 200 Increase 10/300 GL column.

To purify Nb35 and anti-Fab-Nb, the plasmid encoding the nanobody with a PelB periplasmic signal peptide was transformed into *E. coli* BL21 (DE3) cells. The bacteria were cultured in 2L of LB media and grown until the OD600 reached 0.6. Afterward, 200 μM IPTG was added, and the culture was incubated at 18°C with shaking for 16 hours. The bacterial cells were harvested by centrifugation and resuspended in 100 mL of SET buffer (0.5 M sucrose; 0.5 mM EDTA; 0.2 M Tris, pH 8.0). The suspension was stirred at room temperature for 45 minutes. Following this, 200 mL of water was added to the lysate to induce osmotic shock, and the mixture was stirred for an additional 45 minutes. To halt the reaction, final concentration of 150 mM NaCl, 2 mM MgCl_2_, and 20 mM imidazole were added. The lysate was clarified by centrifugation, and the supernatant was loaded onto Ni-NTA resin. The purification process was then continued as described above. After elution from the Ni-NTA resin, the nanobody was further purified using a Superdex 75 Increase 10/300 GL column.

### Expression and purification of GPR3-miniGα_**s**_ fusion protein, GPR3-BRIL and GPR6-BRIL fusion proteins

For the expression of either GPR3 construct, Expi293 cells were transfected with corresponding plasmid using PEI-MAX (Polyethylenimine “Max”, Polysciences) at density of 2.0×10^6^ cell/ml. 44 to 48 hours post-transfection, cells were harvested by centrifugation at 4,000 g for 10 min, rapidly frozen by liquid nitrogen and stored at -80°C until purification.

To purify the GPR3-miniGα_s_ fusion proteins, cells were lysed in hypotonic buffer (25 mM HEPES, pH 7.4, 50 mM NaCl) by a glass dounce homogenizer. The membrane fraction was collected by centrifugation at 40,000 g at 4 °C and solubilized in solubilization buffer (20 mM HEPES pH 7.4, 150 mM NaCl, 10 mM CaCl_2_, 1% LMNG and 0.1% CHS). The homogenate was stirred at 4 °C for 2 h before being centrifugated at 40,000 g for 1 h at 4 °C. Supernatant was filtered through a 0.45 μm membrane filter, and loaded onto M1-Flag agarose beads. The beads were washed sequentially with AP wash buffer I (20 mM HEPES, pH 7.4, 150 mM NaCl, 2 mM CaCl_2_, 0.01% LMNG, 0.001% CHS) and AP wash buffer II (20 mM HEPES, pH 7.4, 150 mM NaCl, 2 mM CaCl_2_, 2 mM ATP, 2 mM KCl, 10 mM MgCl_2_, 0.01% LMNG, 0.001% CHS) followed by a final wash with AP wash buffer I. The bound proteins were eluted with AP elution buffer (20 mM HEPES, pH 7.4, 150 mM NaCl, 5 mM EDTA, 0.1 mg/ml FLAG peptide, 0.01% LMNG, and 0.001% CHS). The eluate was concentrated and purified using a Superose 6 increase 10/300 GL column in SEC buffer (20 mM HEPES, pH 7.4, 150 mM NaCl, 0.01% LMNG, 0.001% CHS) to remove aggregation and peptide.

For the purification of the GPR3-BRIL fusion protein, the membrane fraction was isolated as described above and was solubilized in Solubilization Buffer (20 mM HEPES, 150 mM NaCl, 2 mM CaCl_2_, 0.5 % LMNG, 0.05 % CHS) overnight at 4 °C. The following day, the insoluble material was removed by centrifugation at 180,000 rpm for 40 min. The supernatant was then passed through a 0.45 μm membrane filter, and load onto the M1-Flag agarose beads pre-equilibrated with wash Buffer (20 mM HEPES, 150 mM NaCl, 2 mM CaCl_2_, 0.01 % LMNG, 0.001 % CHS). The beads were washed by wash Buffer until no protein can be detected in the flow-through. GPR3-BIRL protein was eluted by Elution Buffer (20 mM HEPES, 150 mM NaCl, 5 mM EDTA, 0.1 mg/ml flag peptide, 0.01 % LMNG, 0.001 % CHS). Protein was concentrated and further purified by Superose 6 increase 10/300 GL column with SEC Buffer (20 mM HEPES, 150 mM NaCl, 0.01 % LMNG, 0.001 % CHS).

The purification procedure for GPR3-BRIL bound to AF64394 was identical to the above method, except that AF64394 was added at a final concentration of 3 μM throughout the procedure. The purification of GPR6-BRIL protein followed the same procedure as that for GPR3-BRIL.

### Complex assembly

The purified GPR3-miniGα_s_ fusion protein was mixed with Gβ_1_γ_2_ and Nb35 at molar ratios of 1:1.2:1.5, respectively. The mixture was incubated at 4°C for 1 hour, after which the complexes were purified using a Superose 6 Increase 10/300 GL column with SEC buffer. The peak fractions corresponding to the complex were pooled and concentrated to 8-10 mg/mL for cryo-EM sample preparation.

Purified GPR3-BRIL or GPR6-BRIL was mixed with anti-BRIL-Fab and anti-Fab-Nb at molar ratios of 1:1.2:1.5. The complexes were incubated on ice for 1-2 hours, followed by purification using a Superose 6 Increase 10/300 GL column in SEC buffer. The peak fractions were pooled and concentrated to 8-10 mg/mL for cryo-EM sample preparation.

For the assembly of the AF64394-bound GPR3-BRIL complex, the same procedure was followed as described above, with the addition of 3 μM AF64394 throughout the process.

### Cryo-EM sample preparation and data collection

Glow-charged 300 mesh holey carbon grids (Quantifoil Au/Cu R1.2/1.3) were loaded with 2.7-3 μl protein, blotted for 5.5 s in a FEI Vitrobot MarkIV (ThermoFisher Scientific) with the chamber maintained at 8 °C and 100% humidity, and plunge-frozen into liquid ethane. Cryo-EM movies for all complexes were collected on a Titan Krios (Thermo Fisher Scientifc) operated at 300 kV, equipped with a BioQuantum GIF/K3 (Gatan) direct electron detector in a super-resolution mode at a nominal magnification of × 64,000. Data collection were performed using EPU at a dose rate of 22 e^-^ pixel^-1^ s^-1^ for a total dose of 50 e^-^ Å^-2^ over 32 frames. Cryo-EM data collection parameters are summarized in Table EV1.

### Cryo-EM data processing

A total of 3,545 movies were collected for the GPR3-miniGα_s_ complex, 2,630 movies for the GPR3-BRIL complex, and 1,520 movies for the AF64394-bound GPR3-BRIL complex. Movies were motion-corrected and 2×binned to a pixel size of 1.08 Å using MotionCor2 (Zheng, Palovcak et al., 2017). Contrast transfer function (CTF) was estimated using patch-based CTF estimation in cryoSPARC (Punjani, Rubinstein et al., 2017).

For GPR3-miniGα_s_/Gβ_1_γ_2_-C68S/Nb35 complex, particles were initially picked by blob picker and subjected to a round of 2D classification in cryoSPARC. Good 2D class averages of dimeric and monomeric complex were used separately as templates for a second round of particles picking using the template picker. For the dimeric complex, the picked particles underwent two rounds of 2D classification, followed by ab-initio reconstruction and one round of heterogenous refinement. Finally, non-uniform (NU) refinement and local refinement were performed on the best class from heterogenous refinement in cryoSPARC, resulting in a density map with a global resolution of 2.70 Å for the dimeric GPR3-miniGα_s_ structure. For the monomeric complex, particles from good 2D classes after two rounds of 2D classification were further separated through two rounds of 3D classification in RELION (Scheres, 2012). Following this, NU refinement and local refinement in cryoSPARC were performed on the best class from 3D classification to obtain a density map with a global resolution of 3.14 Å.

For the AF64394-bound GPR3-BRIL structure, all data processing was performed in cryoSPARC. Particles were first picked using a round of blob picker and template picker, followed by 2D classification to select clearly defined dimer particles for Topaz model training. The final particles were selected by the Topaz model, and one round of 2D classification and selection was performed. Ab initio reconstruction, heterogeneous refinement, NU refinement, and local refinement were sequentially performed, yielding a final density map with a global resolution of 3.16 Å.

For the GPR3-BRIL apo structure, 1,356 movies collected initially underwent a round of blob picker and template picker, yielding 3,146,401 particles. Particles were also picked from the same movies using the Topaz model, trained using the AF64394-bound GPR3-BRIL structure, resulting in 656,478 particles. The two particle sets were merged, and duplicate particles were removed. The merged particles were then subjected to one round of 2D classification, and dimeric 2D class averages were selected. For the additional 1,274 movies, the Topaz model was used to select 348,568 particles, followed by 2D classification and selection. Afterward, particles from both sets were merged and subjected to ab initio reconstruction and heterogeneous refinement. Finally, NU refinement and local refinement were performed, resulting in a density map of the GPR3-BRIL structure with a global resolution of 3.32 Å. All maps were post-processed using DeepEMhancer (Sanchez-Garcia, Gomez-Blanco et al., 2021).

### Model building

The structure of GPR3, predicted by AlphaFold, and the structure of the miniGα_s_, Gβ_1_γ_2_, and Nb35 complex, derived from our studies (PDB: 8KH5) (Nie et al., 2023), were docked into the cryo-EM density map using Chimera. The model of GPR3-BRIL fusion, generated using AlphaFold, along with structures of ani-BRIL Fab and Nb (derived from PDB: 6WW2) were fit into the map using Chimera. All models were manually adjusted in COOT and refined in Phenix using the secondary structure restraints.

### cAMP accumulation assay

For the cAMP accumulation assay, Expi293F cells were maintained in serum-free SMM 293-TII medium and seeded into six-well plates at a density of 2 × 10^6^ cells/ml. 2 ml cells were co-transfected with 1000 ng of the cAMP biosensor plasmid pGloSensor™-22F (Promega) and 400 ng to 2000 ng of the GPCR plasmids. The amount of GPCR plasmid used for transfection were adjusted to ensure all GPCR mutants have similar surface expression level compared to the WT. 24 hours post-transfection, cells were collected, washed once with Hank’s Balanced Salt Solution (HBSS) and resuspended in cAMP assay buffer (HBSS supplemented with 0.01% bovine serum albumin (BSA, Sigma), 10 mM HEPES (pH 7.3, Beyotime) and 500 μg/ml D-luciferin (Beyotime)). The cell suspension was then seeded in black-bottom 96-well plates with 100 μl per well. After a 30 minutes equilibration period at room temperature, the relative light units (RLUs) for each well were measured using Spark® microplate reader (Tecan). For assays in which GPCR needed to be activated or inhibited by drugs, RLU were read 5 minutes after the addition of 1 μl of drug. Data analysis was performed in GraphPad Prism 7.0 (GraphPad Software). All experiments were performed for at least three times.

### Flow cytometry

Cell surface expression level of the WT and mutant GPCRs was assessed using fluorescence-activated cell sorting (FACS). Expi293F cells (2 mL at a density of 2.0 × 10^6^ cells/mL) were transfected with Flag-tagged GPCR plasmids using PEI-MAX and cultured in six-well plates. After 24 h, 200 μl of the cell suspension was centrifuged at 250 × g for 3 min to remove the culture medium. The resulting cell pellet was washed twice with FACS buffer (HBSS +10 mM HEPES + 2 mM CaCl_2_ + 0.01% BSA, pH 7.4). Cells were stained by Alexa Fluor 647-conjugated M1-Flag antibody (50 μl FACS Buffer + 0.5 μl antibody) and incubated on ice for 15 min. Unbound antibody was removed by an additional wash with FACS buffer. Finally, surface expression levels were quantified by measuring the fluorescence using a BD Accuri™ C6 Plus flow cytometer.

### Fluorescence-detection size exclusion chromatography (FSEC)

2ml of Expi293F cells were transfected with a GPCR-mCherry fusion construct by PEI-MAX. After 24 h, cells were collected by centrifugation and solubilized in 200 μl Solubilization Buffer (20 mM HEPES, 150 mM NaCl, 100 μM TCEP, 1 % DDM, 0.1 % CHS) for 2 hours at 4 °C with rotation. The lysis was then centrifuged at 30,000 g for 10 min to remove insoluble debris. The supernatant was loaded onto a gel filtration column (Zenix-C SEC-300 7.8/300 300 Å, Sepax Technologies, Inc.) in high-performance liquid chromatograph (HPLC), using a buffer containing 20 mM HEPES, 150 mM NaCl, 100 μM TCEP, 0.03 % DDM, 0.003 % CHS.

## Data availability

The atomic structures have been deposited at the Protein Data Bank (PDB) under the accession codes XX. The EM maps have been deposited at the Electron Microscopy Data Bank (EMDB) under the accession numbers XX.

## Acknowledgements

We thank staff at Shuimu BioSciences for their help with cryo-EM data collection.

## Author contributions

Z.Q and Y.N. purified the GPR3-BRIL fusion complex, collected cryo-EM data and performed cryo-EM data processing and model building under the supervision of S.Z. W.W purified the GPR3-miniGs fusion complex, collected EM data and built model under the supervision of S.Z. Z.Q, W.W and Y.N., performed cAMP accumulation assay with assistance from J.L, B.Z and H.X. S.Z, Z.Q, W.W and Y.N. wrote the manuscripts.

## Competing interests

The authors declare no competing interests.

## Notes

### Competing Interest Statement

The authors have declared no competing interest.

## Reference

Asher WB, Geggier P, Holsey MD, Gilmore GT, Pati AK, Meszaros J, Terry DS, Mathiasen S, Kaliszewski MJ, McCauley MD, Govindaraju A, Zhou Z, Harikumar KG, Jaqaman K, Miller LJ, Smith AW, Blanchard SC, Javitch JA (2021) Single-molecule FRET imaging of GPCR dimers in living cells. Nat Methods 18: 397–405

Barekatain M, Johansson LC, Lam JH, Chang H, Sadybekov AV, Han GW, Russo J, Bliesath J, Brice NL, Carlton MBL, Saikatendu KS, Sun H, Murphy ST, Monenschein H, Schiffer HH, Popov P, Lutomski CA, Robinson CV, Liu ZJ, Hua T et al. (2024) Structural insights into the high basal activity and inverse agonism of the orphan receptor GPR6 implicated in Parkinson’s disease. Sci Signal 17: eado8741

Bouvier M (2001) Oligomerization of G-protein-coupled transmitter receptors. Nat Rev Neurosci 2: 274–86

Carpenter B, Tate CG (2016) Engineering a minimal G protein to facilitate crystallisation of G protein-coupled receptors in their active conformation. Protein Eng Des Sel 29: 583–594

Chabre M, le Maire M (2005) Monomeric G-protein-coupled receptor as a functional unit. Biochemistry 44: 9395–403

Chen G, Staffen N, Wu Z, Xu X, Pan J, Inoue A, Shi T, Gmeiner P, Du Y, Xu J (2024) Structural and functional characterization of the endogenous agonist for orphan receptor GPR3. Cell Res 34: 262–265

Dong T, Hu G, Fan Z, Wang H, Gao Y, Wang S, Xu H, Yaffe MB, Vander Heiden MG, Lv G, Chen J (2024) Activation of GPR3-beta-arrestin2-PKM2 pathway in Kupffer cells stimulates glycolysis and inhibits obesity and liver pathogenesis. Nat Commun 15: 807

Eggerickx D, Denef JF, Labbe O, Hayashi Y, Refetoff S, Vassart G, Parmentier M, Libert F (1995) Molecular cloning of an orphan G-protein-coupled receptor that constitutively activates adenylate cyclase. Biochem J 309 (Pt 3): 837–43

Farran B (2017) An update on the physiological and therapeutic relevance of GPCR oligomers. Pharmacol Res 117: 303–327

Fotiadis D, Jastrzebska B, Philippsen A, Muller DJ, Palczewski K, Engel A (2006) Structure of the rhodopsin dimer: a working model for G-protein-coupled receptors. Curr Opin Struct Biol 16: 252–9

Freudzon L, Norris RP, Hand AR, Tanaka S, Saeki Y, Jones TL, Rasenick MM, Berlot CH, Mehlmann LM, Jaffe LA (2005) Regulation of meiotic prophase arrest in mouse oocytes by GPR3, a constitutive activator of the Gs G protein. J Cell Biol 171: 255–65

Godlewski G, Jourdan T, Szanda G, Tam J, Cinar R, Harvey-White J, Liu J, Mukhopadhyay B, Pacher P, Ming Mo F, Osei-Hyiaman D, Kunos G (2015) Mice lacking GPR3 receptors display late-onset obese phenotype due to impaired thermogenic function in brown adipose tissue. Sci Rep 5: 14953

Gurevich VV, Gurevich EV (2018) GPCRs and Signal Transducers: Interaction Stoichiometry. Trends Pharmacol Sci 39: 672–684

Gusach A, Garcia-Nafria J, Tate CG (2023) New insights into GPCR coupling and dimerisation from cryo-EM structures. Curr Opin Struct Biol 80: 102574

Hinckley M, Vaccari S, Horner K, Chen R, Conti M (2005) The G-protein-coupled receptors GPR3 and GPR12 are involved in cAMP signaling and maintenance of meiotic arrest in rodent oocytes. Dev Biol 287: 249–61

Huang Y, Rafael Guimaraes T, Todd N, Ferguson C, Weiss KM, Stauffer FR, McDermott B, Hurtle BT, Saito T, Saido TC, MacDonald ML, Homanics GE, Thathiah A (2022) G protein-biased GPR3 signaling ameliorates amyloid pathology in a preclinical Alzheimer’s disease mouse model. Proc Natl Acad Sci U S A 119: e2204828119

Huang Y, Skwarek-Maruszewska A, Horre K, Vandewyer E, Wolfs L, Snellinx A, Saito T, Radaelli E, Corthout N, Colombelli J, Lo AC, Van Aerschot L, Callaerts-Vegh Z, Trabzuni D, Bossers K, Verhaagen J, Ryten M, Munck S, D’Hooge R, Swaab DF et al. (2015) Loss of GPR3 reduces the amyloid plaque burden and improves memory in Alzheimer’s disease mouse models. Sci Transl Med 7: 309ra164

Ignatov A, Lintzel J, Hermans-Borgmeyer I, Kreienkamp HJ, Joost P, Thomsen S, Methner A, Schaller HC (2003) Role of the G-protein-coupled receptor GPR12 as high-affinity receptor for sphingosylphosphorylcholine and its expression and function in brain development. J Neurosci 23: 907–14

Inoue A, Ishiguro J, Kitamura H, Arima N, Okutani M, Shuto A, Higashiyama S, Ohwada T, Arai H, Makide K, Aoki J (2012) TGFalpha shedding assay: an accurate and versatile method for detecting GPCR activation. Nat Methods 9: 1021–9

Kolakowski LF, Jr. (1994) GCRDb: a G-protein-coupled receptor database. Recept Channels 2: 1–7

Kroeze WK, Sassano MF, Huang XP, Lansu K, McCorvy JD, Giguere PM, Sciaky N, Roth BL (2015) PRESTO-Tango as an open-source resource for interrogation of the druggable human GPCRome. Nat Struct Mol Biol 22: 362–9

Ledent C, Demeestere I, Blum D, Petermans J, Hamalainen T, Smits G, Vassart G (2005) Premature ovarian aging in mice deficient for Gpr3. Proc Natl Acad Sci U S A 102: 8922–6

Lin S, Han S, Cai X, Tan Q, Zhou K, Wang D, Wang X, Du J, Yi C, Chu X, Dai A, Zhou Y, Chen Y, Zhou Y, Liu H, Liu J, Yang D, Wang MW, Zhao Q, Wu B (2021) Structures of G(i)-bound metabotropic glutamate receptors mGlu2 and mGlu4. Nature 594: 583–588

Liu X, Masoudi A, Kahsai AW, Huang LY, Pani B, Staus DP, Shim PJ, Hirata K, Simhal RK, Schwalb AM, Rambarat PK, Ahn S, Lefkowitz RJ, Kobilka B (2019) Mechanism of beta2AR regulation by an intracellular positive allosteric modulator. Science 364: 1283–1287

Lorente JS, Sokolov AV, Ferguson G, Schioth HB, Hauser AS, Gloriam DE (2025) GPCR drug discovery: new agents, targets and indications. Nat Rev Drug Discov

Lu S, Jang W, Inoue A, Lambert NA (2021) Constitutive G protein coupling profiles of understudied orphan GPCRs. PLoS One 16: e0247743

Mehlmann LM, Saeki Y, Tanaka S, Brennan TJ, Evsikov AV, Pendola FL, Knowles BB, Eppig JJ, Jaffe LA (2004) The Gs-linked receptor GPR3 maintains meiotic arrest in mammalian oocytes. Science 306: 1947–50

Milligan G, Ward RJ, Marsango S (2019) GPCR homo-oligomerization. Curr Opin Cell Biol 57: 40–47

Nie Y, Qiu Z, Chen S, Chen Z, Song X, Ma Y, Huang N, Cyster JG, Zheng S (2023) Specific binding of GPR174 by endogenous lysophosphatidylserine leads to high constitutive G(s) signaling. Nat Commun 14: 5901

Oeckl P, Hengerer B, Ferger B (2014) G-protein coupled receptor 6 deficiency alters striatal dopamine and cAMP concentrations and reduces dyskinesia in a mouse model of Parkinson’s disease. Exp Neurol 257: 1–9

Papasergi-Scott MM, Perez-Hernandez G, Batebi H, Gao Y, Eskici G, Seven AB, Panova O, Hilger D, Casiraghi M, He F, Maul L, Gmeiner P, Kobilka BK, Hildebrand PW, Skiniotis G (2024) Time-resolved cryo-EM of G-protein activation by a GPCR. Nature 629: 1182–1191

Punjani A, Rubinstein JL, Fleet DJ, Brubaker MA (2017) cryoSPARC: algorithms for rapid unsupervised cryo-EM structure determination. Nat Methods 14: 290–296

Rasmussen SG, DeVree BT, Zou Y, Kruse AC, Chung KY, Kobilka TS, Thian FS, Chae PS, Pardon E, Calinski D, Mathiesen JM, Shah ST, Lyons JA, Caffrey M, Gellman SH, Steyaert J, Skiniotis G, Weis WI, Sunahara RK, Kobilka BK (2011) Crystal structure of the beta2 adrenergic receptor-Gs protein complex. Nature 477: 549–55

Ruiz-Medina J, Ledent C, Valverde O (2011) GPR3 orphan receptor is involved in neuropathic pain after peripheral nerve injury and regulates morphine-induced antinociception. Neuropharmacology 61: 43–50

Russell IC, Zhang X, Bumbak F, McNeill SM, Josephs TM, Leeming MG, Christopoulos G, Venugopal H, Flocco MM, Sexton PM, Wootten D, Belousoff MJ (2024) Lipid-Dependent Activation of the Orphan G Protein-Coupled Receptor, GPR3. Biochemistry 63: 625–631

Sanchez-Garcia R, Gomez-Blanco J, Cuervo A, Carazo JM, Sorzano COS, Vargas J (2021) DeepEMhancer: a deep learning solution for cryo-EM volume post-processing. Commun Biol 4: 874

Saotome K, McGoldrick LL, Ho JH, Ramlall TF, Shah S, Moore MJ, Kim JH, Leidich R, Olson WC, Franklin MC (2025) Structural insights into CXCR4 modulation and oligomerization. Nat Struct Mol Biol 32: 315–325

Scheres SH (2012) RELION: implementation of a Bayesian approach to cryo-EM structure determination. J Struct Biol 180: 519–30

Shen C, Mao C, Xu C, Jin N, Zhang H, Shen DD, Shen Q, Wang X, Hou T, Chen Z, Rondard P, Pin JP, Zhang Y, Liu J (2021) Structural basis of GABA(B) receptor-G(i) protein coupling. Nature 594: 594–598

Sveidahl Johansen O, Ma T, Hansen JB, Markussen LK, Schreiber R, Reverte-Salisa L, Dong H, Christensen DP, Sun W, Gnad T, Karavaeva I, Nielsen TS, Kooijman S, Cero C, Dmytriyeva O, Shen Y, Razzoli M, O’Brien SL, Kuipers EN, Nielsen CH et al. (2021) Lipolysis drives expression of the constitutively active receptor GPR3 to induce adipose thermogenesis. Cell 184: 3502–3518 e33

Tanaka S, Ishii K, Kasai K, Yoon SO, Saeki Y (2007) Neural expression of G protein-coupled receptors GPR3, GPR6, and GPR12 up-regulates cyclic AMP levels and promotes neurite outgrowth. J Biol Chem 282: 10506–15

Tanaka S, Shimada N, Shiraki H, Miyagi T, Harada K, Hide I, Sakai N (2022) GPR3 accelerates neurite outgrowth and neuronal polarity formation via PI3 kinase-mediating signaling pathway in cultured primary neurons. Mol Cell Neurosci 118: 103691

Teng X, Chen S, Nie Y, Xiao P, Yu X, Shao Z, Zheng S (2022a) Ligand recognition and biased agonism of the D1 dopamine receptor. Nat Commun 13: 3186

Teng X, Chen S, Wang Q, Chen Z, Wang X, Huang N, Zheng S (2022b) Structural insights into G protein activation by D1 dopamine receptor. Sci Adv 8: eabo4158

Thathiah A, Spittaels K, Hoffmann M, Staes M, Cohen A, Horre K, Vanbrabant M, Coun F, Baekelandt V, Delacourte A, Fischer DF, Pollet D, De Strooper B, Merchiers P (2009) The orphan G protein-coupled receptor 3 modulates amyloid-beta peptide generation in neurons. Science 323: 946–51

Tsutsumi N, Mukherjee S, Waghray D, Janda CY, Jude KM, Miao Y, Burg JS, Aduri NG, Kossiakoff AA, Gati C, Garcia KC (2020) Structure of human Frizzled5 by fiducial-assisted cryo-EM supports a heterodimeric mechanism of canonical Wnt signaling. Elife 9

Uhlenbrock K, Gassenhuber H, Kostenis E (2002) Sphingosine 1-phosphate is a ligand of the human gpr3, gpr6 and gpr12 family of constitutively active G protein-coupled receptors. Cell Signal 14: 941–53

Valverde O, Celerier E, Baranyi M, Vanderhaeghen P, Maldonado R, Sperlagh B, Vassart G, Ledent C (2009) GPR3 receptor, a novel actor in the emotional-like responses. PLoS One 4: e4704

Velazhahan V, Ma N, Pandy-Szekeres G, Kooistra AJ, Lee Y, Gloriam DE, Vaidehi N, Tate CG (2021) Structure of the class D GPCR Ste2 dimer coupled to two G proteins. Nature 589: 148–153

Xiong Y, Xu Z, Li X, Wang Y, Zhao J, Wang N, Duan Y, Xia R, Han Z, Qian Y, Liang J, Zhang A, Guo C, Inoue A, Xia Y, Chen Z, He Y (2024) Identification of oleic acid as an endogenous ligand of GPR3. Cell Res 34: 232–244

Yue Y, Liu L, Wu LJ, Wu Y, Wang L, Li F, Liu J, Han GW, Chen B, Lin X, Brouillette RL, Breault E, Longpre JM, Shi S, Lei H, Sarret P, Stevens RC, Hanson MA, Xu F (2022) Structural insight into apelin receptor-G protein stoichiometry. Nat Struct Mol Biol 29: 688–697

Zhang K, Wu H, Hoppe N, Manglik A, Cheng Y (2022) Fusion protein strategies for cryo-EM study of G protein-coupled receptors. Nat Commun 13: 4366

Zheng SQ, Palovcak E, Armache JP, Verba KA, Cheng Y, Agard DA (2017) MotionCor2: anisotropic correction of beam-induced motion for improved cryo-electron microscopy. Nat Methods 14: 331–332

Zhuang Y, Krumm B, Zhang H, Zhou XE, Wang Y, Huang XP, Liu Y, Cheng X, Jiang Y, Jiang H, Zhang C, Yi W, Roth BL, Zhang Y, Xu HE (2021) Mechanism of dopamine binding and allosteric modulation of the human D1 dopamine receptor. Cell Res

